# Effects of cover crops on beetle assemblages in tea plantations

**DOI:** 10.1101/2021.03.19.436210

**Authors:** Li-Lin Chen, Gabor Pozsgai, Xiang-Yu Li, Lei Li, Gadi V.P. Reddy, Min-Sheng You

**Author notes:** Correspondence:* Min-Sheng You, State Key Laboratory of Ecological Pest Control for Fujian and Taiwan Crops, College of Plant Protection, Fuzhou, Fujian 350002, China., Gabor Pozsgai, State Key Laboratory of Ecological Pest Control for Fujian and Taiwan Crops, College of Plant Protection, Fuzhou, Fujian 350002, China.

## Abstract

Beetles are visible members of food webs in tea plantations, with high species richness and abundance. Many tea pests, as well as natural enemies, are members of this order, so a knowledge of how groundcovers affect beetles can aid pest management. We collected beetles in a replicated field experiment in the Wuyi Mountains, Fujian Province China. Tea was intercropped with *Paspalum notatum* or *Chamaecrista rotundifolia,* or rows were cleared to bare ground, or in the control they were left unmanaged to allow weeds to grow naturally. Sampling, done by sweep netting and vegetation beating, was conducted monthly, between May 2006 and April 2008, and Coleoptera abundance, biomass, species richness and assemblage structures were compared between groundcover treatments. Total beetle abundance and species richness were significantly higher in tea intercropped with *C. rotundifolia* and bare ground than in naturally grown weedy control. Whilst there was no difference between predator assemblages among treatments for any measure, herbivores were more abundant, weighed more, and were more diverse in *C. rotundifolia* treatments than in weedy control. Biomass and species richness were also greater in plots with *P. notatum* groundcover than those in weedy control. We found that beetle assemblages varied both seasonally and with ground cover treatment, but the potential pest control impact of more species-rich beetle assemblages was mixed, and further work is needed to gain information on trophic groups with potential benefits for use in non-insecticidal pest management.

## Introduction

Tea (*Camellia sinensis* Kuntze, 1887) is a perennial, evergreen shrub that is widely cultivated throughout the tropics and subtropics, particularly in hilly or mountainous regions (Hazarika *et al.*, 2009; Zhang *et al.*, 2019). Pest damage to tea, however, can reduce its marketability and cause significant losses in yield (Muraleedharan & Chen, 1997). Such problems have led to the overuse of pesticides, which in turn can contaminate tea with pesticide residues, cause pest resistance, and induce pest resurgence due to destruction of natural enemies (Fernandez *et al.*, 1993; Sarkar & Hajra, 1994). Tea production is increasingly associated with prolonged and extensive use of synthetic pesticides. Such use, along with the expansion of crop monocultures, has resulted in pest outbreaks in tea, likely because of the adverse effects of those two management practices on predators and parasitoids (Sudoi *et al.*, 1996; Wang *et al.*, 2004). Tea is, however, an important export commodity, and the use of pesticides on this cash crop has recently become more carefully regulated by several agencies and organizations (Ford, 2018; Chen HP *et al.*, 2019; Juncker, 2019). Also, interest has increased in alternative control strategies to replace or reduce use of pesticides and lower their resulting residues (Bian *et al.*, 2014; Guo *et al.*, 2019). Most of these alternative practices focus on efforts to enhance biocontrol ecosystem services provided by natural enemies (Chen *et al.*, 2020).

The beetles (Coleoptera) are an abundant and species-rich order that includes both pests and beneficial species important in tea production. Herbivorous beetles, including bark borers, leaf eating species, and shoot hole borers, can severely reduce tea yield and quality (Table 1). Predatory beetles are believed to help suppress pest Hemiptera in tea (Table 1).

**Table 1.**
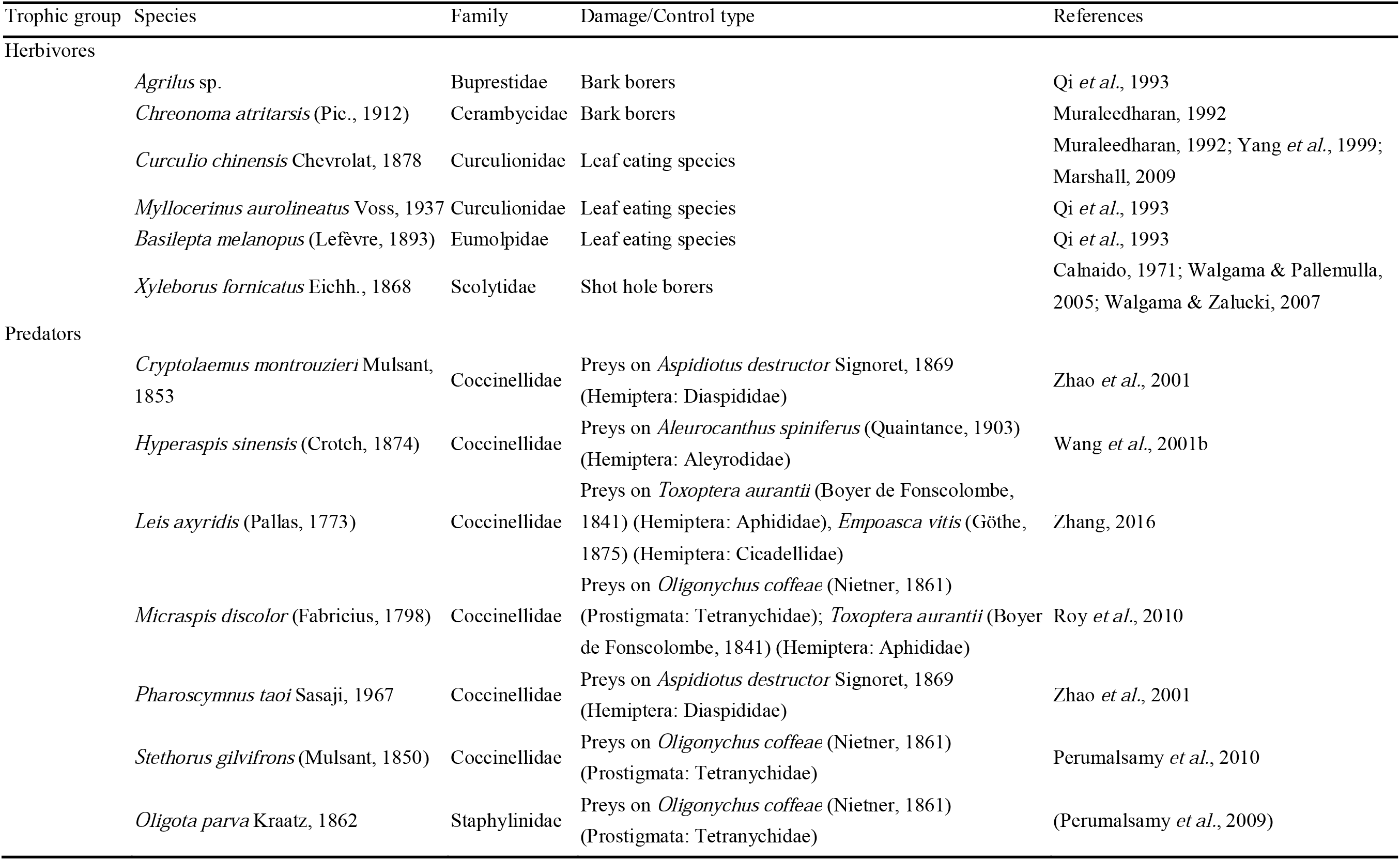
Effects of some Coleoptera species in tea plantations.

Since the densities of both pests and their natural enemies are influenced by the surrounding vegetation, it is reasonable to manage this environment to enhance biocontrol services if possible (Bastola *et al.*, 2016; Li *et al.*, 2016a; Gurr *et al.*, 2016, 2017).

Indeed, intercropping with a variety of cover crops has been used in many crop systems to enhance the action of beneficial organisms. Cover crops can be used to reduce soil erosion, improve soil structure or fertility, increase water infiltration, suppress weeds or nematodes, and improve crop quality (Armecin *et al.*, 2005; Song *et al.*, 2006; Wu *et al.*, 2013; Schipanski et al. 2014a). Cover crops that are properly selected and well managed can enhance the abundance and efficacy of natural enemies (Bugg *et al.*, 1991; Landis *et al.*, 2000; Zhao *et al.*, 2016), mostly by providing them with essential resources, such as pollen, honeydew, and nectar (Gurr & Nicol, 2000; Gurr *et al.*, 2017), and providing habitat for mating, oviposition, or shelter (Hossain *et al.*, 2002; Pappas *et al.*, 2017).

Existing evidence on efficacy of cover crops in enhancing beneficial species in tea plantations is mixed (Chen *et al.*, 2019a, b). Song et al. (2006) found that predatory Coleoptera abundance was significantly higher in tea interplanted with white clover (*Trifolium repens* L., 1753) compared to tea monocultures, and this change led to better suppression of hemipteran pests. Several plants, including *Corymbia citriodora* (Hook.) Hill et Johnson, 1848 and *Lavandula pinnata* Lundmark, 1780 (Zhang *et al.*, 2014a), *Senna tora* (L.) Roxb. (Zhang *et al.*, 2014b), and *Hedyotis uncinella* Hook. et Arn., 1833 have also been successfully employed as tea cover crops (Chen, 2002; Song *et al.*, 2006; Zhang *et al.*, 2014a, b).

However, it tea, there is little information on the effect of cover crops on groups like beetles, which can act both as pests and natural enemies, or how cover crops might affect the pest: natural enemy ratios. The impacts of particular cover crop species on beetle assemblages is also poorly understood. Our goal was to assess the effects of cover crops on assemblages of beetles in tea canopies. We tested the hypothesis that ground covers in tea plantations influence the abundance, biomass, and species richness of both herbivore and predatory beetles, affecting the whole assemblage. We predicted that intercropping in tea would increase natural enemy: pest ratios compared to a naturally weedy control, and that this effect would be equally observable for abundance, biomass, and species richness.

## Materials and methods

### Study sites

Field experiments were conducted between May 2006 and April 2008 in a 7-year-old Oolong tea (clone Wuyi Shuixian) plantation located at Xingcun in the Wuyi Mountains, Fujian Province, China (27°38′51.4′′ N and 117°54′29.9′′ E, 736 m altitude). The tea plantation is situated on red soil and the regional climate is classified as humid subtropical monsoon.

### Experimental design

Four ground cover treatments - (1) *sown Paspalum notatum* Flügge, 1810 (Poaceae) (Pas), (2) sown *Chamaecrista rotundifolia* Greene, 1899 (Fabaceae) (Cas), (3) bare ground (Bar), and (4) natural weedy ground cover (Nat) - were compared in a fully randomized design with three replicates of each treatment. Naturally grown weedy ground cover served as the control in our experiment. *P. notatum* and *C. rotundifolia* were manually sown in May 2006 at a seed rate of 45 kg/ha and 7.5 kg/ha, respectively. In the bare ground treatment weeds were manually cleared each month without the use of herbicide. Tea intercropped with *P. notatum* or *C. rotundifolia* were kept nearly weed-free by mowing, and both treatments and natural ground cover were mowed between tea rows to a 5 cm stubble height in both years in September. Plot size was 340 m^2^ (17 m × 20 m), and plots were spaced 5 m apart. Experimental plots differed from the conventionally managed areas only by the intercropping strategy employed and freedom from the use of insecticides. In all other respects they were maintained following the recommended agronomic practice and had similar natural conditions, such as geographical features, topography, and soil texture.

### Insect sampling

The tea canopy was sampled monthly. Samples were taken from five randomly selected 1 m^2^ areas of canopy in each plot. Six white trays (36 cm × 45 cm) were placed beneath tea bushes, 30 sweeps with a net (38.1 cm diameter hoop) were performed, and then the tea canopy was beaten 20 times. Insects in nets and on trays were transferred to plastic bags and frozen for later identification to species or morphospecies. Average wet weight of the insects preserved in alcohol was measured individually for 1 to 80 specimens of each collected species to the nearest mg with a Swiss Mettler AB204-S electronic balance for insect biomass estimation. Before being weighed, specimens were dried on absorbent paper to remove excess alcohol (as per Zintzen *et al.*, 2008). For each species, the average weight (n = 1-80, depending on availability) was taken as the biomass value for that species. This biomass value was then multiplied by the number of individuals of that species in each sample.

Beetles were assigned to trophic groups as herbivores, predators, omnivores, or saprophages, based on the literature (Pang & Mao, 1979; Zhang & Tan, 2004; Zhang & Li, 2011; Ren *et al.*, 2016) and behavioral observations at the study sites. Voucher specimens were preserved in 75% alcohol and stored at the Institute of Applied Ecology, Fujian Agriculture and Forestry University.

### Data analysis

Data from the five subsamples were pooled, and mean species richness, mean abundance, and mean biomass of the replicates were calculated for total Coleoptera and for each of the four trophic groups, respectively. Because of their limited importance compared to predators and herbivores, these values for saprophagous and omnivorous trophic groups were not analyzed separately, and species belonging to these groups were only included in the analysis when overall numbers were compared.

Linear mixed effect models (LMER) were used to compare beetle richness, abundance and biomass amongst treatments with sampling months included as a fixed effect and sampling dates as a random effect. Satterthwaite’s method for calculating degrees of freedom was used to obtain P-values, with the help of the *lmerTest* R package (Kuznetsova *et al.*, 2017). Least-squares means were computed, and all treatment combinations compared using the *lsmeans* package (Lenth, 2016) in R (R Development Core Team, 2019) and P-values adjusted using Tukey’s method. All comparisons were repeated separately for all beetles, herbivores only, predatory beetles only, and for the predator: herbivore ratios. The normality of both within-cluster residuals and the residuals of random effects were individually checked using standard Q-Q plots and Shapiro-tests. Homogeneity of the variances was tested using Levene-tests.

Whether or not treatments had a significant effect on beetle assemblages was tested using principal response curve analysis (PRC), with repeated observations. Permutational ANOVA with 999 iterations was used to validate the model, and the most influential species were highlighted. PRC was repeated for pooled beetle abundance and biomass, herbivore and predator abundances each separately, and herbivore and predator biomass each separately. Multivariate analysis was conducted using the *vegan* (Oksanen *et al.*, 2019) R package.

For LMER models both count and biomass data were log-transformed, as log(*x*+1) and log(*x*+0.01), respectively. Before calculating the predator: herbivore ratios, count data were log(*x*+1), and biomass data were cube-root 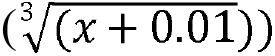 transformed, and the cube-root and log-transformed values of these proportions, respectively, were included in the model as response variables. Richness proportions were cube-root transformed. For multivariate analysis, abundance data were transformed using Hellinger transformation and biomass data were log-transformed (log(*x*+0.01)).

## Results

### Overview of results

Altogether, 28,394 individuals of 117 species of beetles were captured during the 23 months of this experiment (Table 2.). Of these 117 species, 66 were categorized as herbivores, 29 as predators, and 22 as other feeding types that included omnivores, saprophages, or mixed feeders (see Supplementary material 1 for the full species list and categorization).

**Table 2.** Abundance, biomass, and species richness of beetles collected in intercropped tea plantations, under four different groundcover treatments, in China over a two year period, segregated by beetle feeding type.

|  | Feeding trait | Herbivore | Omnivore | Predator | Saprophage | Total |
| --- | --- | --- | --- | --- | --- | --- |
| Abundance | Pas<sup>†</sup> | 5341 | 21 | 1334 | 162 | 6858 |
|  | Cas<sup>‡</sup> | 5893 | 23 | 1290 | 251 | 7457 |
|  | Bar<sup>§</sup> | 6841 | 22 | 940 | 163 | 7966 |
|  | Nat<sup>††</sup> | 5196 | 11 | 795 | 111 | 6113 |
|  | Total | 23271 | 77 | 4359 | 687 | 28394 |
| Biomass | Pas | 252.43 | 0.16 | 2.42 | 0.33 | 255.35 |
|  | Cas | 215.66 | 0.16 | 2.49 | 0.40 | 218.70 |
|  | Bar | 292.50 | 0.18 | 1.61 | 0.30 | 294.59 |
|  | Nat | 245.94 | 0.09 | 1.08 | 0.16 | 247.27 |
|  | Total | 1006.54 | 0.58 | 7.60 | 1.19 | 1015.91 |
| Species richness | Pas | 41 | 6 | 16 | 5 | 68 |
|  | Cas | 33 | 6 | 19 | 5 | 63 |
|  | Bar | 34 | 10 | 17 | 4 | 65 |
|  | Nat | 36 | 6 | 20 | 4 | 66 |
|  | Total | 66 | 16 | 29 | 6 | 117 |

<sup>†</sup> Pas: sown *Paspalum notatum*; <sup>‡</sup> Cas: sown *Chamaecrista rotundifolia*; <sup>§</sup> Bar: bare ground; <sup>††</sup> Nat: natural ground cover.

### Abundance

Both treatment and sampling month significantly affected differences between the log-transformed beetle abundance but their combined effect was not significant. This model had a total explanatory power of 84.9%, in which the fixed effects explained 67.1% of the variance (Table 3). Pairwise comparison of the treatments revealed significantly greater log-abundance in Cas treatment than in Nat (*P* = 0.014, Fig. 1 A). There was no significant effect of treatment for herbivorous beetle, but treatment did have a significant effect for predator log-abundance (Table 3).

**Table 3.** Results of REML modelling for log abundance, log biomass, and species richness, separately for pooled data, for feeding traits, and for herbivore-predator ratios. Total and fixed effect (in brackets) variances explained by the model are indicated.

|  | *Sum Sq* | *Mean Sq* | *Num Df* | *Den Df* | *Statistic* | *P-Value* |
| --- | --- | --- | --- | --- | --- | --- |
| *Pooled log-abundance Variance explained (%) = 84.9 (67.1)* |  |  |  |  |  |  |
| *TR* | 2.816 | 0.939 | 3 | 217 | 3.641 | 0.014* |
| *month* | 20.285 | 1.844 | 11 | 11 | 7.153 | 0.001** |
| *TR:month* | 4.610 | 0.140 | 33 | 217 | 0.542 | 0.981 |
| *Hervibore log-abundance Variance explained (%) = 88.8 (69.9)* |  |  |  |  |  |  |
| *TR* | 1.808 | 0.603 | 3 | 217 | 1.452 | 0.229 |
| *month* | 32.735 | 2.976 | 11 | 11 | 7.171 | 0.001** |
| *TR:month* | 13.596 | 0.412 | 33 | 217 | 0.993 | 0.485 |
| *Predator log-abundance Variance explained (%) = 71.0 (27.7)* |  |  |  |  |  |  |
| *TR* | 7.011 | 2.337 | 3 | 217 | 7.012 | < 0.001*** |
| *month* | 3.758 | 0.342 | 11 | 11 | 1.025 | 0.484 |
| *TR: month* | 10.060 | 0.305 | 33 | 217 | 0.915 | 0.606 |
| *Predator-herbivore log-abundance ratios Variance explained (%) = 30.5 (19.5)* |  |  |  |  |  |  |
| *TR* | 0.353 | 0.118 | 3 | 217 | 0.766 | 0.514 |
| *month* | 3.477 | 0.316 | 11 | 11 | 2.054 | 0.124 |
| *TR:month* | 5.637 | 0.171 | 33 | 217 | 1.110 | 0.321 |
| *Pooled log-biomass Variance explained (%) = 87.9 (46.8)* |  |  |  |  |  |  |
| *TR* | 1.567 | 0.522 | 3 | 217 | 0.714 | 0.545 |
| *month* | 18.102 | 1.646 | 11 | 11 | 2.249 | 0.097 |
| *TR:month* | 19.477 | 0.590 | 33 | 217 | 0.807 | 0.766 |
| *Herbivore log-biomass Variance explained (%) = 86.6 (46.8)* |  |  |  |  |  |  |
| *TR* | 1.433 | 0.478 | 3 | 217 | 0.498 | 0.684 |
| *month* | 24.233 | 2.203 | 11 | 11 | 2.296 | 0.092 |
| *TR:month* | 31.360 | 0.950 | 33 | 217 | 0.991 | 0.488 |
| *Predator log-biomass Variance explained (%) = 60.6 (25.0)* |  |  |  |  |  |  |
| *TR* | 2.883 | 0.961 | 3 | 217 | 4.205 | 0.006** |
| *month* | 2.463 | 0.224 | 11 | 11 | 0.980 | 0.513 |
| *TR:month* | 8.405 | 0.255 | 33 | 217 | 1.114 | 0.316 |
| *Predator-herbivore log-biomass ratios Variance explained (%) = 69.4 (40.3)* |  |  |  |  |  |  |
| *term* |  |  |  |  |  |  |
| *TR* | 0.271 | 0.090 | 3 | 217 | 0.672 | 0.570 |
| *month* | 4.392 | 0.399 | 11 | 11 | 2.972 | 0.042* |
| *TR:month* | 4.554 | 0.138 | 33 | 217 | 1.027 | 0.434 |
| *Pooled species richness Variance explained (%) = 55.7 (20.4)* |  |  |  |  |  |  |
| *TR* | 53.493 | 17.831 | 3 | 217 | 4.822 | 0.003** |
| *month* | 28.270 | 2.570 | 11 | 11 | 0.695 | 0.722 |
| *TR:month* | 113.293 | 3.433 | 33 | 217 | 0.928 | 0.584 |
| *Herbivore species richness Variance explained (%) = 52.4 (11.4)* |  |  |  |  |  |  |
| *TR* | 4.116 | 1.372 | 3 | 217 | 1.005 | 0.391 |
| *month* | 3.406 | 0.310 | 11 | 11 | 0.227 | 0.990 |
| *TR:month* | 46.565 | 1.411 | 33 | 217 | 1.034 | 0.424 |
| *Predator species richness Variance explained (%) = 49.8 (28.6)* |  |  |  |  |  |  |
| *TR* | 14.531 | 4.844 | 3 | 217 | 5.001 | 0.002** |
| *month* | 15.452 | 1.405 | 11 | 11 | 1.450 | 0.274 |
| *TR:month* | 42.616 | 1.291 | 33 | 217 | 1.333 | 0.117 |
| *Predator-herbivore species richness ratios Variance explained (%) = 46.1 (16.7)* |  |  |  |  |  |  |
| *TR* | 0.044 | 0.015 | 3 | 217 | 0.544 | 0.653 |
| *month* | 0.177 | 0.016 | 11 | 11 | 0.600 | 0.795 |
| *TR:month* | 0.975 | 0.030 | 33 | 217 | 1.099 | 0.335 |

Significance codes: ‘\*’, $P < 0.05$ ; ‘\*\*’, $P < 0.01$ ; ‘\*\*\*’, $P < 0.001$ .

**Fig. 1.**
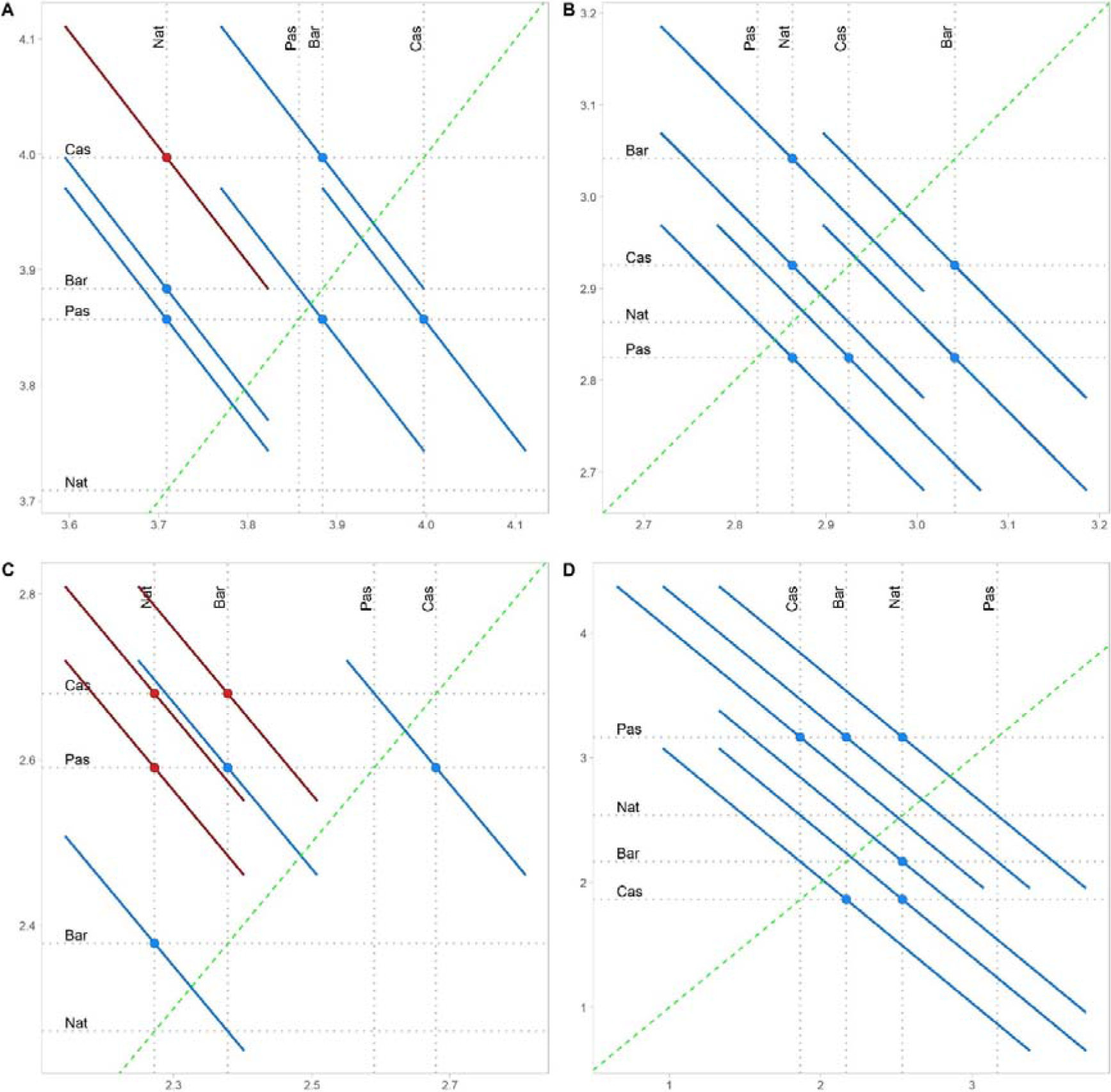
Diffograms showing the differences amongst the calculated least-square means of treatments for A) total log-abundance, B) herbivore log-abundance, C) predator log-abundance, and the D) predator:herbivore log-abundance ratios. In each individual graph, the green, dashed, diagonal line indicates zero difference between two least-square means (line of equality), and each solid line represent one pairwise comparison between treatments. When pairwise comparison lines do not cross the line of equality, differences are significant (*P* < 0.05 after adjusting with the Tukey’s method). Significant differences are indicated with red color, lines with non-significant differences are blue. Horizontal and vertical axes are identical, indicating the range of least-square means. Pas: sown *Paspalum notatum*; Cas: sown *Chamaecrista rotundifolia*; Bar: bare ground; Nat: natural ground cover.

Log-abundance of the predators differed significantly between the Nat and Cas groundcovers (P < 0.001), the Nat and Pas groundcovers (P = 0.009), and the Bar and Cas ground covers (P = 0.015), with predators being less abundant in Nat than in Pas or Cas, and more abundant in Cas than in Nat (Fig. 1 B-C). There was no significant difference between the treatments in predator–herbivore log-abundance ratios (Table 3, Fig. 1 D). Detailed time series comparisons between treatment effects on abundance in relation to sampling date are shown in Supplementary material 2 and Supplementary Material 3.

### Biomass

Although the model based on total biomass explained 87.89% of the variance, only 46.79% of this amount was attributed to the fixed effects and neither sampling month nor treatments had a statistically significant relationship with total beetle biomass (Table 3, Fig. 2 A). Similarly, none of the fixed variables showed a significant effect when only herbivores were included in the model (Table 3). Predator biomass, on the other hand, was significantly different between treatments (Table 3, Fig. 2 B), with both intercropping treatments having greater beetle biomass than the Nat treatment (Pas: *P* = 0.02, Cas: *P* = 0.01, Fig. 2 C). The ratios of predators to herbivores did not show a significant treatment effect, but the effect of sampling month was significant (Table 3). For detailed information on model outputs readers should consult Supplementary material 2 and Supplementary material 4 for monthly comparison of treatments on biomass.

**Fig. 2.**
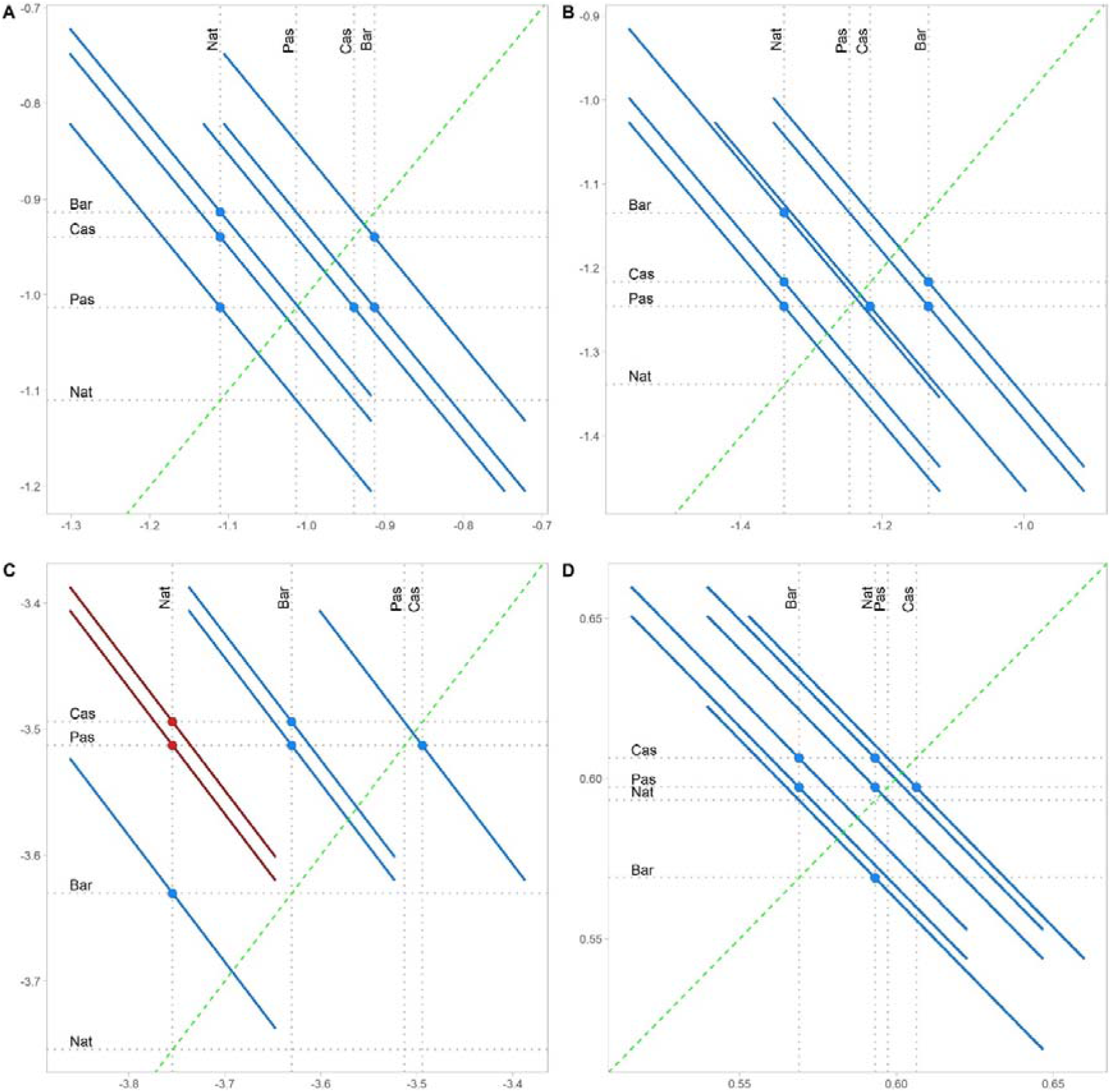
Diffograms showing the differences amongst the calculated least-square means of treatments for **A)** total log-biomass, **B)** herbivore log-biomass, **C)** predator log-biomass, and the **D)** predator: herbivore log-biomass ratios. In each individual graph diagonal green, dashed line indicates zero difference between two least-square means (line of equality), and each solid line represent one pairwise comparison between treatments. When pairwise comparison lines do not cross the line of equality differences are significant (*P* < 0.05 after adjusting with the Tukey’s method). Significant differences are indicated with red color, lines with non-significant differences are blue. Horizontal and vertical axes are identical, indicating the range of least-square means. Pas: sown *Paspalum notatum*; Cas: sown *Chamaecrista rotundifolia*; Bar: bare ground; Nat: natural ground cover.

### Species richness

Total species richness was significantly greater in Bar and Cas treatments compared to the Nat treatment (*P* = 0.014 and *P* = 0.004, respectively), while marginally significantly greater in the Pas treatment (*P* = 0.051). This model explains 55.74% of the total variance, of which the fixed effects explain 20.42% (Table 3, Fig. 3 A).

**Fig. 3.**
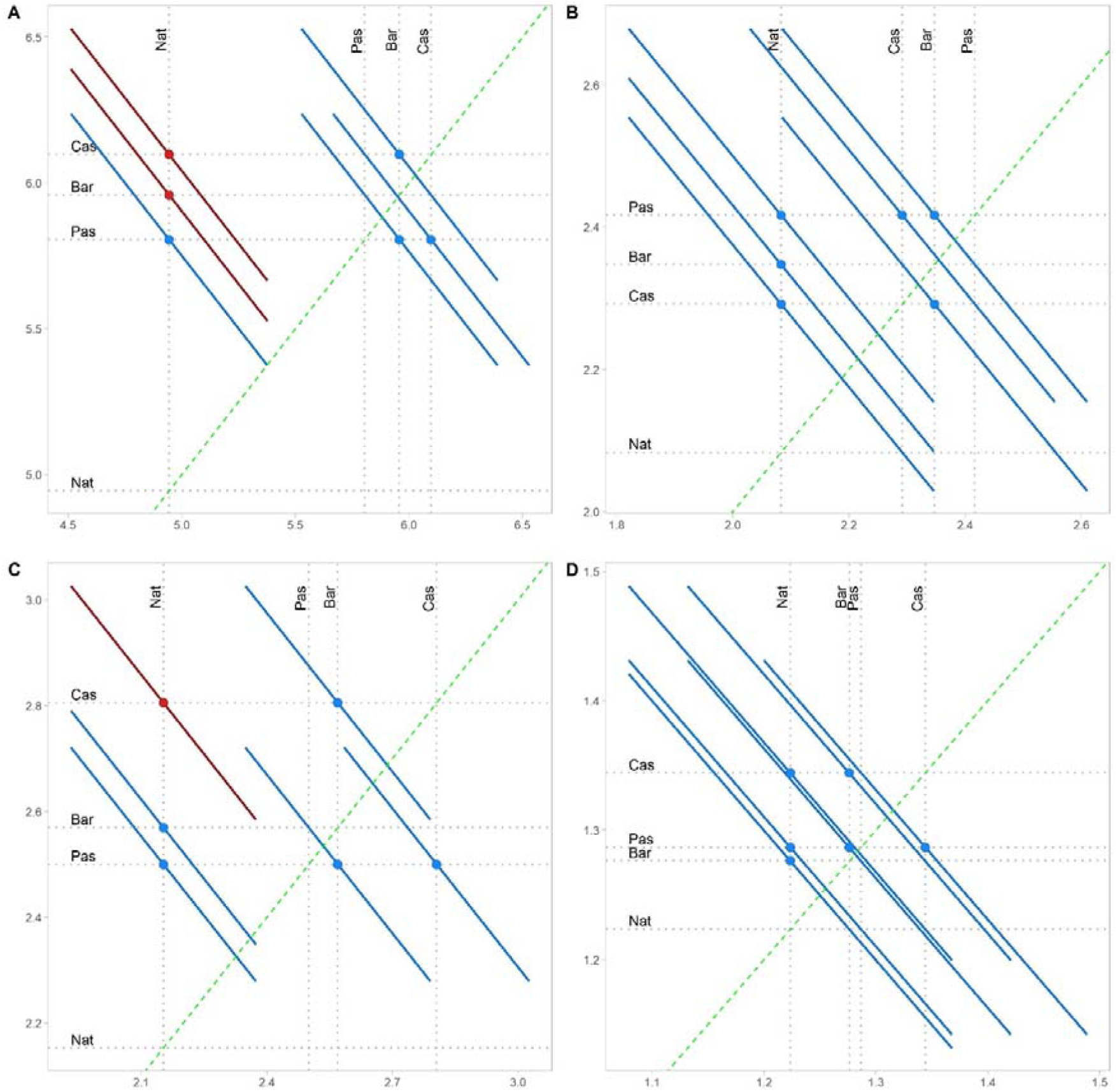
Diffograms showing the differences amongst the calculated least-square means of treatments for **A)** total species richness, **B)** herbivore species richness, **C)** predator species richness, and the **D)** predator: herbivore species richness ratios. In each individual graph, diagonal green dashed line indicates zero difference between two least-square means (line of equality), and each solid line represent one pairwise comparison between treatments. When pairwise comparison lines do not cross the line of equality differences are significant (*P* < 0.05 after adjusting with the Tukey’s method). Significant differences are indicated with red color, lines with non-significant differences are blue. Horizontal and vertical axes are identical indicating the range of least-square means. Pas: sown *Paspalum notatum*; Cas: sown *Chamaecrista rotundifolia*; Bar: bare ground; Nat: natural ground cover.

None of the fixed effects indicated any significant difference when only herbivore richness was considered but treatment effect was significant for predators alone (Table 3), showing greater species richness in both Bar and Cas treatments than in the Nas treatment (*P* = 0.073 and *P* = 0.001, respectively, Fig. 3 B-C). The total explanatory power for the predatory species richness model was 49.85%, of which the fixed effects explained 28.60%. Detailed numerical outputs of the model can be found in Supplementary material 2 and time-series plots of treatment effects on species richness-related variables are shown in Supplementary material 5.

### Treatment Effect on Beetle Assemblages

Principal response curve analysis showed significant treatment effect for species assemblages when 1) total species abundance, 2) only herbivore abundance, and 3) predator biomass variables were used as the input data matrix (Table 4).

**Table 4.** Results of permutational ANOVAs on principal response curve (PRC) models, based on different community matrices. Number of permutations is 999, and degrees of freedom is 69 in every cases. Statistically significant models are highlighted with bold font face.

| Assemblage based on | Variance | F-Value | P-Value |
| --- | --- | --- | --- |
| Pooled abundance | 0.063 | 1.11 | 0.014* |
| Herbivore abundance | 0.087 | 1.19 | 0.002** |
| Predator abundance | 0.078 | 1.06 | 0.200 |
| Pooled biomass | 0.755 | 1.08 | 0.180 |
| Herbivore biomass | 0.009 | 0.64 | 0.990 |
| Predator biomass | < 0.001 | 1.19 | 0.048* |

Significance codes: ‘\*’, $P < 0.05$ ; ‘\*\*’, $P < 0.01$

Effects show a distinctive seasonality for abundance-related datasets, whereas this temporal influence was not apparent when biomass-based datasets were used. There was an obvious dominance of herbivores in the assemblages, regardless of whether data matrices were based on abundance or biomass. The most abundant species - *Epuraea luteola* Erichson, 1843 (Nitidulidae), *Migneauxia lederi* Reitter, 1875 (Lathriidae), *Myllocerinus aurolineatus* Voss, 1937 (Curculionidae), and *Basilepta melanopus* (Lefèvre, 1893) (Eumolpidae) - become more dominant over time within the treatments, while *Serangium japonicum* Chapin, 1940 (Coccinellidae) and *Pharoscymnus taoi* Sasaji, 1967 (Coccinellidae) decreased in relative importance over time. However, if only predators were included in the model, only *P. taoi* was affected negatively. This decline in relative importance in the assemblages was only prominent when models were based on abundance. Compared to the control (Nas), biomass-based importance of *Cryptogonus postimedialis* Kapur, 1948 (Coccinellidae) increased in all treatments (Fig. 4).

**Fig. 4.**
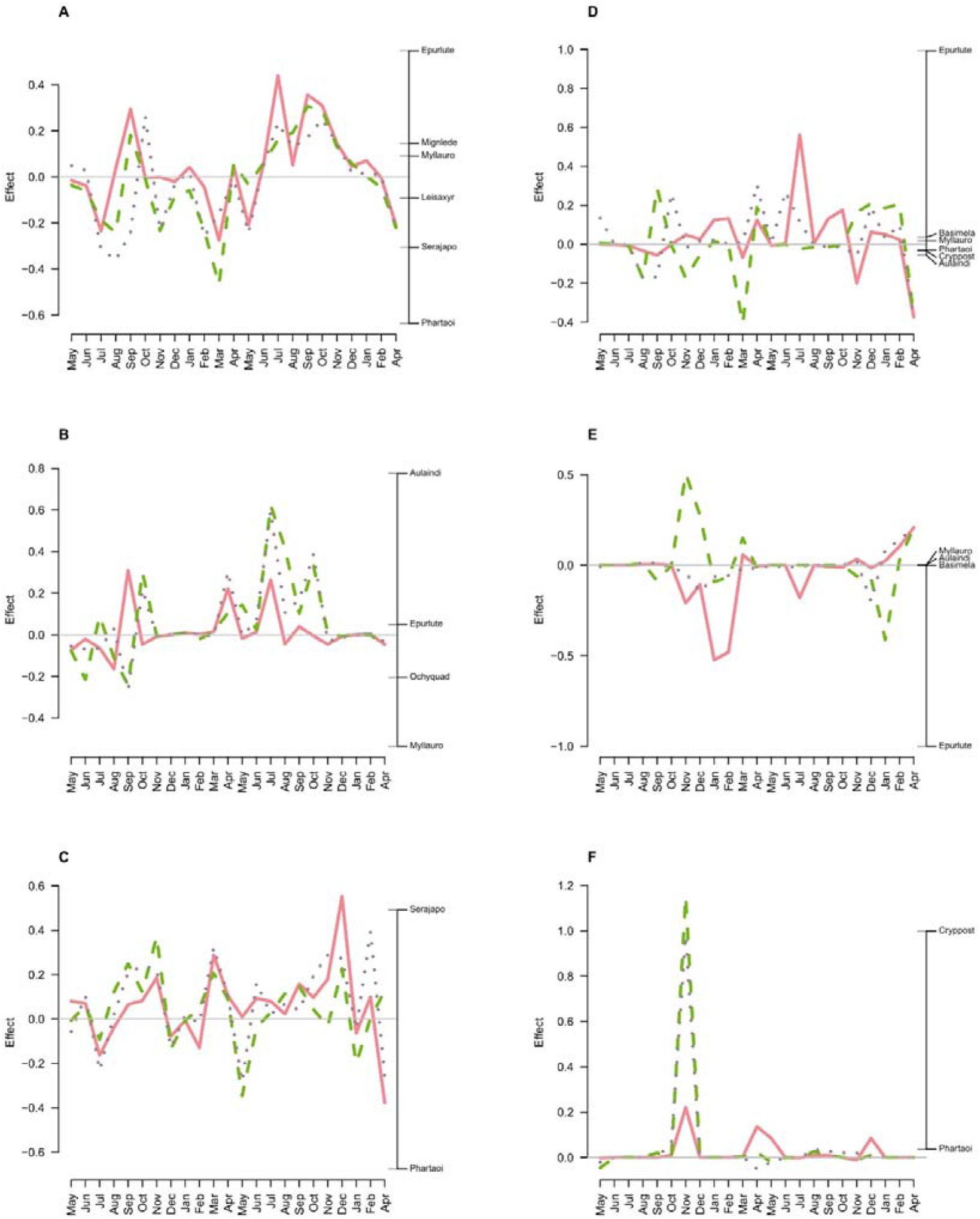
Principal response curves models based on **A)** abundance data when total species are included, **B)** abundance data of herbivore species, **C)** abundance data of predatory species, **D)** biomass data of total species, **E)** biomass data of herbivore species, and **F)** biomass data of predatory species. Time range starts in May, 2006, and it ends in April 2008. The solid red line represents the bare ground treatment, the dashed green line, the *Chamaecrista rotundifolia* treatment and the dotted green-brown line, the *Paspalum notatum* treatment. The horizontal grey line indicates the natural ground cover control. The most influential species in shaping assemblage differences are marked on the right side of the graph. Abbreviations as: Aulaindi - *Aulacophora indica*, Basimela - *Basilepta melanopus*, Cryppost - *Cryptogonus postmedialis*, Epurlute –*Epuraea luteola*, Leisaxyr - *Leis axyridis*, Mignlede - *Migneauxia lederi*, Myllauro - *Myllocerinus aurolineatus*, Ochyquad - *Ochyromera quadrimaculata*, Phartaoi - *Pharoscymnus taoi*, Serajapo - *Serangium japonicum*.

## Discussion

In this study we investigated the effects of cover crops on beetle assemblages in tea plantations and showed distinctive, yet sometimes weak, diversity increases with cover crop treatments. We mostly found species that are common in tea growing areas, with a high herbivore dominance in the assemblages (Marshall, 1938; Benjamin, 1968; Shiao, 1977; Qi *et al.*, 1993; Wang *et al.*, 2001a, Zhao *et al.*, 2001; Chen *et al.*, 2009a; Dai, 2010).

We found greater abundance and species richness in the Cas treatment than in the Nat control, indicating a positive effect of this cover crops on beetle biodiversity. Cover crops are generally thought to provide a more sheltered, favorable overwintering microclimate and offer more food than bare ground (Altieri *et al.*, 1985; Lassau *et al.*, 2005; Gurr *et al.*, 2017). In our case, however, although *C. rotundifolia* appeared to attract a greater beetle biodiversity into the tea canopy than naturally weedy rows, there was no significant difference between the two sown treatments and plots where tea rows were manually weeded (Bar). Whereas this seemingly contradicts the improved microhabitat hypothesis, the explanation may lie in the seasonality of our treatments. Cover crops were mowed for the winter; thus, the sudden disturbance and the decreased amount of spatially diverse overwintering habitats made the refuge potential of all four treatments similar.

Indeed, there was a distinct seasonality observable in our dataset: both the total abundance and total biomass increased substantially in the winter months (December to February) (Ye *et al.*, 2010; Chen, 2011), while species richness remained within the previously observed range (Supplementary material 3-5). This increase was mainly caused by the large numbers of the nitidulid *E. luteola*. Since this winter effect was similar in all treatments, it did not result in a statistically significant treatment effect. The role of the evergreen tea as a winter refuge for herbivorous insects may be the driving factor in shaping this phenomenon, but the fact that there was no comparable growth in numbers of predatory beetles, which could have used the same habitat as refuge, does not support this hypothesis. Alternatively, the timing of tea flowering in Wuyi coincided with the mass appearance of a pollen feeding insect in this period, which may offer a better explanation (Chen *et al.*, 2009b). Indeed, the predator: herbivore ratios decreased during these months, pointing to a disproportional increase in pollen feeders.

Both predatory beetle abundance and biomass were greater in both of the two sown treatments compared to the control, but this did not translate to a reduction in the number of herbivores. Whether this was the result of an insufficient increase of effective predators to control herbivores, or merely seasonality or other variables masking the effect is yet to be clarified. Indeed, the positive effect of cover crops on the abundance and biomass of natural enemies was more visible in the first year, and it became less pronounced in the second year. This change suggests that temporal factors, particularly the growing period of the cover crop, played an important role in determining to what extent a given plant species, or species mix, can contribute toward increasing natural enemy abundance (Bugg *et al.*, 1991; Wei *et al.*, 2010; Schipanski *et al.*, 2014b; Trichard *et al.*, 2014; Rivers et al. 2020). Alternatively, since flowering leguminous plants such as *C. rotundifolia* can support a greater abundance of herbivorous Coleoptera, particularly *E. luteola* and various weevils, and these plants also produce dense foliage that is favorable for herbivores (Chen, 2011), the net effect on increased predator numbers may be nil because herbivores have also increased.

Another potential explanation of why in some months, abundance differences between treatments could not been detected, could be that in intercropped tea gardens, beetles did not move onto tea plants if a suitable cover vegetation was available. Consequently, it is unclear if cover crops, which provide abundant resources, would act as a “source” or as a “sink” for predatory beetles in agroecosystems (Corbett & Plant, 1993).

Nonetheless, large populations of a few species of predators may be insufficient to efficiently suppress a wide variety of pests (Bugg *et al.*, 1991). Moreover, greater predator abundance may not necessarily improve pest control, particularly of herbivorous beetles, when predatory beetles do not particularly target the key pest species (Bugg *et al.*, 1991; Baggen & Gurr, 1998). Increasing biodiversity allows a more efficient resource exploitation (Snyder *et al.*, 2006; Li *et al.*, 2016b), and consequently, the increase in species richness (as a proxy for biodiversity) in our Cas treatments implies a potential ability of this cover crop to result in pest suppression.

Both in terms of total biodiversity and abundance or richness of predatory beetles, Cas generally performed better than Pas. Nevertheless, weeded, bare ground rows (Bar) also showed significantly higher species richness than in the control. The underlying mechanisms of the two significant differences, however, are probably different; *C. rotundifolia* likely attracts beneficial insects by providing shelter or other resources (Chen *et al.*, 2019a, b), whereas bare soil surfaces open up opportunities for ambush predators to easily find prey (Schmidt & Rypstra, 2010). Since these normally generalist species that are specifically adapted to disturbed habitats are more likely to stay in the adjacent field margins, their long-term value in tea canopies as biological control agents may be limited.

On the other hand, one of the additional benefits of cover crops may be promoting biodiversity including the diversity of economically insignificant species (Nicholls & Altieri 2013; Li *et al.*, 2016; de Pedro *et al.*, 2020), and thus they can be of high conservation interest. Theoretically, since cover crops are a more heterogeneous ecosystem with available niches (Rosenzweig, 1995; Dufour *et al.*, 2006), they should support more species. However, for long-term increase of arthropod biodiversity, ephemeral habitats like seasonal cover crops, are less likely to be beneficial than more stable habitats (Rivers *et al.*, 2020). Therefore, if nature conservation is also to be considered, choosing the right cover plant(s) is crucial.

Whilst treatments in our study influenced beetle assemblages, temporal fluctuations of the superdominant species *E. luteola* made the effects inconclusive. Despite the significant differences in predator abundance, abundance-based assemblages of predators did not show significant treatment effects. Alternately, fluctuating populations of two of the, most likely competing, coccinellids (*S. japonicum* and *P. taoi*) was also visible, and this may have been what was driving the changes in these beetle assemblages.

While the results of this study illustrate the influence of cover crops on beetle assemblages in tea plantations, all of our treatments had advantages and disadvantages. If there are no obvious economic benefits of sown cover crops over naturally occurring weed cover (i.e., no increase from cover crops in herbivore control), then farmers are likely to choose the least labor–intensive, yet profitable management (Song *et al.*, 2006; Zhan *et al.*, 2018; Chen *et al.*, 2019a; Rivers *et al.*, 2020). Since natural ground cover is unlikely to be favored by farmers (even though it may host considerable biodiversity), planting a variety of cover crops may be an acceptable compromise between entirely cleared and weedy rows in tea plantations. Besides promoting biodiversity, with a careful selection of plant species mix the additional benefits (such as soil physical and chemical properties improvement, carbon retention, microbial activity enhancement, and erosion control) of plant cover would be maximized (Wu *et al.*, 2013; Sahoo *et al.*, 2016; Fu *et al.*, 2018; Khan & Verma, 2018; Zhan *et al.*, 2018; Fu et al. 2020). This approach would still provide the benefits of a diverse cover vegetation, and as a consequence of greater diversity of other organisms, which would likely reduce overwintering pest herbivores in tea plantations. Assessing the detailed effects of plant species, or any mixture of these, from a wide ecological perspective is needed though before practitioners and farmers can be advised of their value. Further research is necessary to understand other aspects, particularly seasonality and temporal variation, of how cover crops drive beetle diversity and assemblages. Simultaneously maximizing biodiversity and pest control using cover crops may be a copious task but the delivery of these multiple ecosystem services will benefit both sustainable agriculture and nature conservation.

## Acknowledgments

We express our appreciation to the Planters and Managers of Fujian Wuyishan Yongsheng Industrial Tea Company Ltd. for providing us with a tea plantation for our field experiments in the Wuyi Mountains, Fujian Province, China. We thank Jenni A. Stockan for her edits and correction of the English in the final version of the manuscript. This work was supported by the National Natural Science Foundation of China (No. 31501650), National Key R & D Program of China (No. 2016YFD0200900), Fujian Agriculture and Forestry University Construction Project for Technological Innovation and Service System of Tea Industry Chain (No. K1520005 A03), Fujian Spark Program (No. 2018S0019), “111” Program: Innovation Center for Ecologically Based Pest Management of Subtropical Crops (No. D16012).

## Supplementary information

**Supplementary material 1** List of full species names, families they belong to, their feeding guild categorization, and abbreviations used.

**Supplementary material 2** All model statistics and pairwise least-square means comparisons from the repeated measures linear mixed effect models (LMER). Pas: sown *Paspalum notatum*; Cas: sown *Chamaecrista rotundifolia*; Bar: bare ground; Nat: natural ground cover.

**Supplementary material 3** Boxplot comparisons of abundance-related measures for each sampling date. (A) shows total log-abundance, (B) herbivore log-abundance, (C) predator log-abundance, and (D) predator:herbivore log-abundance ratios Pas: sown *Paspalum notatum*; Cas: sown *Chamaecrista rotundifolia*; Bar: bare ground; Nat: natural ground cover.

**Supplementary material 4** Boxplot comparisons of biomass-related measures for each sampling date. (A) shows total log-biomass, (B) herbivore log-biomass, (C) predator log-biomass, and (D) predator-herbivore log-biomass ratios. Pas: sown *Paspalum notatum*; Cas: sown *Chamaecrista rotundifolia*; Bar: bare ground; Nat: natural ground cover.

**Supplementary material 5** Boxplot comparisons of richness-related measures for each sampling date. (A) shows total species richness, (B) herbivore species richness, (C) predator species richness, and (D) predator:herbivore species richness ratios. Pas: sown *Paspalum notatum*; Cas: sown *Chamaecrista rotundifolia*; Bar: bare ground; Nat: natural ground cover.

