## Supplementary Material 1 for "Effects of cover crops on beetle assemblages in tea plantations"

| Family | Species | Feeding guild | | Abundance | | Biomass |
| --- | --- | --- | --- | --- | --- | --- |
| Anthicidae | Anthicidae sp. | Herbivore | | 13 | | 0.0155 |
|  | *Omonadus floralis* (L., 1758) | Herbivore | | 1 | | 0.0018 |
| Apionidae | *Apion* sp. | Herbivore | | 9 | | 0.0630 |
| Bruchidae | *Spermophagus rufiventris* (Boheman, 1833) | Herbivore | | 4 | | 0.0013 |
|  | *Zabrotes subfasciatus* (Boheman, 1833) | Herbivore | | 1 | | 0.0009 |
| Buprestidae | *Agrilus* sp. | Herbivore | | 9 | | 0.0279 |
| Carabidae | *Stenolophus agonoides* Bates, 1883 | Herbivore | | 7 | | 0.0810 |
|  | *Stenolophus connotatus* Bates, 1873 | Herbivore | | 3 | | 0.0681 |
|  | *Stenolophus difficilis* (Hope, 1845) | Herbivore | | 4 | | 0.0072 |
| Chrysomelidae | *Agelastica coerulea* Baly, 1874 | Herbivore | | 1 | | 0.0020 |
|  | *Aphthona strigosa* Baly, 1874 | Herbivore | | 45 | | 0.0290 |
|  | *Arthrotus nigrofasciatus* Jacoby, 1890 | Herbivore | | 41 | | 0.2744 |
|  | *Aulacophora indica* (Gmelin, 1790) | Herbivore | | 103 | | 2.0301 |
|  | *Chrysolina aurichalcea* (Mannerheim, 1825) | Herbivore | | 1 | | 0.0015 |
|  | *Chrysolina fricata* (Wollaston, 1854) | Herbivore | | 1 | | 0.0158 |
|  | Chrysomelidae sp. | Herbivore | | 2 | | 0.0026 |
|  | *Clitea fulva* Chen, 1933 | Herbivore | | 3 | | 0.0008 |
|  | *Clitea metallica* Chen, 1933 | Herbivore | | 1 | | 0.0042 |
|  | *Dercetina flavocincta* (Hope, 1831) | Herbivore | | 1 | | 0.0019 |
|  | *Gastrolina tonkinea* Chen, 1931 | Herbivore | | 1 | | 0.0011 |
|  | *Longitarsus cyanipennis* Bryant, 1924 | Herbivore | | 3 | | 0.0035 |
|  | *Macrima cornuta* (Laboissiere, 1936) | Herbivore | | 8 | | 0.0632 |
|  | *Monolepta quadriguttata* (Motschulsky, 1860) | Herbivore | | 8 | | 0.0422 |
|  | *Nisotra gemella* (Erichson, 1843) | Herbivore | | 1 | | 0.0028 |
|  | *Oulema atrosuturalis* (Pic, 1923) | Herbivore | | 1 | | 0.0018 |
|  | *Pagria signata* (Motschulsky, 1858) | Herbivore | | 143 | | 0.1330 |
|  | *Phaedon brassicae* Baly, 1874 | Herbivore | | 12 | | 0.0912 |
|  | *Phyllotreta rectilineata* Chen, 1939 | Herbivore | | 18 | | 0.0171 |
|  | *Phyllotreta striolata* (Fabricius, 1801) | Herbivore | | 4 | | 0.0043 |
|  | *Syneta adamsi* Baly, 1877 | Herbivore | | 4 | | 0.0132 |
| Coccinellidae | *Henosepilachna operculata* (Liu, 1950) | Herbivore | | 1 | | 0.0282 |
|  | *Henosepilachna vigintioctopunctata* (Fabricius, 1775) | Herbivore | | 3 | | 0.0126 |
| Crioceridae | *Macroplea japana* Jacoby, 1885 | Herbivore | | 1 | | 0.0010 |
| Curculionidae | *Callosobruchus chinensis* (L., 1758) | Herbivore | | 2 | | 0.0014 |
|  | *Hypera basalis* Voss, 1937 | Herbivore | | 2 | | 0.0216 |
|  | *Hypomeces pulviger* (Herbst, 1795) | Herbivore | | 3 | | 0.0267 |
|  | *Larinus ovalis* Roelofs, 1873 | Herbivore | | 2 | | 0.0184 |
|  | *Myllocerinus aurolineatus* Voss, 1937 | Herbivore | | 1120 | | 13.7041 |
|  | *Nothomyllocerus pelidnus* (Voss, 1958) | Herbivore | | 1 | | 0.0059 |
|  | *Ochyromera quadrimaculata* Voss, 1953 | Herbivore | | 76 | | 0.6709 |
|  | *Phytoscaphus ciliatus* Roelofs, 1873 | Herbivore | | 36 | | 0.0392 |
|  | *Sympiezomias velatus* (Chevrolat, 1845) | Herbivore | | 10 | | 0.0934 |
| Dryophthoridae | *Sitophilus oryzae* Schoenherr, 1838 | Herbivore | | 1 | | 0.0026 |
| Elateridae | *Aeoloderma brachmana* (Candeze, 1859) | Herbivore | | 1 | | 0.0034 |
|  | *Agriotes sericatus* Schwarz, 1891 | Herbivore | | 2 | | 0.0005 |
|  | *Xanthopenthes granulipennis* (Miwa, 1929) | Herbivore | | 3 | | 0.2001 |
| Eumolpidae | *Aulexis atripennis* Pic, 1923 | Herbivore | | 1 | | 0.0059 |
|  | *Aulexis tuberculata* Tan, 1992 | Herbivore | | 9 | | 0.0288 |
|  | *Basilepta melanopus* (Lefèvre, 1893) | Herbivore | | 1308 | | 2.0089 |
|  | *Basilepta ruficollis* (Jacoby, 1885) | Herbivore | | 1 | | 0.0078 |
|  | *Basilepta sinara* (Weise, 1922) | Herbivore | | 4 | | 0.0145 |
|  | *Chiridopsis bipunctata* (L., 1767) | Herbivore | | 23 | | 0.0360 |
|  | *Cleoporus variabilis* (Baly, 1874) | Herbivore | | 1 | | 0.0001 |
|  | *Colasposoma dauricum* Mannerheim, 1849 | Herbivore | | 3 | | 0.1203 |
|  | *Nodina striopunctata* Tan, 1988 | Herbivore | | 3 | | 0.0150 |
|  | *Smaragdina nigrifrons* (Hope, 1842) | Herbivore | | 1 | | 0.0295 |
| Meloidae | *Epicauta obscurocephala* Reitter, 1905 | Herbivore | | 1 | | 0.0024 |
| Melolonthidae | *Apogonia cribricollis* Burmeister, 1855 | Herbivore | | 1 | | 0.0344 |
| Mordellidae | *Mordellistena pumila* (Gyllenhal, 1810) | Herbivore | | 2 | | 0.0046 |
| Scolytidae | *Dryocoetiops coffeae* Schedl, 1964 | Herbivore | | 3 | | 0.0018 |
|  | *Scolytoplatypus superciliosus* Tsai *et* Huang, 1965 | Herbivore | | 1 | | 0.0007 |
| Rutelidae | *Anomala corpulenta* Motschulsky, 1854 | Herbivore | | 1 | | 0.1781 |
| Nitidulidae | *Carpophilus dimidiatus* (Fabricius, 1792) | Herbivore | | 1 | | 0.0011 |
|  | *Epuraea luteola* Erichson, 1843 | **Herbivore** | | 20180 | | 986.7068 |
| Tenebrionidae | *Cteniopinus hypocrita* (Marseul, 1876) | Herbivore | | 7 | | 0.3406 |
|  | *Micropedinus pallidipennis* Lewis, 1894 | Herbivore | | 3 | | 0.0018 |
| Carabidae | *Colliuris* sp. | Predator | | 1 | | 0.0058 |
|  | *Pentagonica subcordicollis* Bates, 1873 | Predator | | 1 | | 0.0135 |
| Coccinellidae | *Aspidimerus ruficrus* Gorham, 1895 | Predator | | 1 | | 0.0051 |
|  | *Cheilomenes sexmaculata* (Fabricius, 1781) | Predator | | 2 | | 0.0616 |
|  | *Chilocorus kuwanae* Silvestri, 1909 | Predator | | 8 | | 0.0864 |
|  | Coccinellidae sp. 1 | Predator | | 10 | | 0.0026 |
|  | Coccinellidae sp. 2 | Predator | | 37 | | 0.0295 |
|  | Coccinellidae sp. 3 | Predator | | 5 | | 0.0512 |
|  | Coccinellidae sp. 4 | Predator | | 14 | | 0.0513 |
|  | *Cryptogonus postimedialis* Kapur, 1948 | Predator | | 322 | | 2.8175 |
|  | *Cryptolaemus montrouzieri* Mulsant, 1853 | Predator | | 4 | | 0.0561 |
|  | *Hyperaspis sinensis* (Crotch, 1874) | Predator | | 2 | | 0.0468 |
|  | *Leis axyridis* (Pallas, 1773) | Predator | | 229 | | 0.2950 |
|  | *Micraspis discolor* (Fabricius, 1798) | Predator | | 1 | | 0.0085 |
|  | *Pharoscymnus taoi* Sasaji, 1967 | Predator | | 2616 | | 2.4351 |
|  | *Platynaspis maculosa* (Weise, 1910) | Predator | | 4 | | 0.0164 |
|  | *Propylea japonica* (Thunberg, 1781) | Predator | | 148 | | 0.7234 |
|  | *Rodolia pumila* Weise, 1892 | Predator | | 1 | | 0.0094 |
|  | *Scymnus babai* Sasaji, 1971 | Predator | | 2 | | 0.0015 |
|  | *Scymnus schmidti* Fürsch, 1958 | Predator | | 33 | | 0.0383 |
|  | *Scymnus yamato* Kamiya, 1961 | Predator | | 2 | | 0.0022 |
|  | *Serangium japonicum* Chapin, 1940 | Predator | | 627 | | 0.4661 |
|  | *Stethorus aptus* Kapur, 1948 | Predator | | 3 | | 0.0072 |
|  | *Stethorus cantonensis* Pang, 1966 | Predator | | 128 | | 0.0282 |
|  | *Stethorus longisiphonulus* Pang, 1966 | Predator | | 1 | | 0.0012 |
|  | *Stethorus Parapauperculus* Pang, 1966 | Predator | | 123 | | 0.3075 |
|  | *Telsimia emarginata* Chapin | Predator | | 2 | | 0.0013 |
| Staphylinidae | *Stenus calliceps* Bernhauer, 1916 | Predator | | 31 | | 0.0264 |
|  | *Stenus cirrus* Benick, 1940 | Predator | | 1 | | 0.0045 |
| Carabidae | *Calleida lepida* Redtenbacher, 1867 | Omnivore | | 3 | | 0.0300 |
|  | *Chlaenius junceus* Andrewes, 1923 | Omnivore | | 1 | | 0.0536 |
|  | *Chlaenius leucops* (Wiedemann, 1823) | Omnivore | | 4 | | 0.0675 |
|  | *Colliuris* sp. | Omnivore | | 23 | | 0.1487 |
|  | *Drypta lineola* MacLeay, 1825 | Omnivore | | 1 | | 0.0130 |
|  | *Euplynes batesi* Harold, 1877 | Omnivore | | 4 | | 0.0040 |
|  | *Harpalus chalcentus* Bates, 1873 | Omnivore | | 1 | | 0.0230 |
|  | *Harpalus sinicus* Hope, 1845 | Omnivore | | 1 | | 0.0543 |
|  | *Harpalus tridens* Morawitz, 1862 | Omnivore | | 2 | | 0.0115 |
|  | *Hexagonia cyclops* (Matsumura, 1910) | Omnivore | | 1 | | 0.0026 |
|  | *Lebia iolanthe* Bates, 1883 | Omnivore | | 2 | | 0.0057 |
|  | *Mastax poecila* Schaum, 1863 | Omnivore | | 1 | | 0.0069 |
| Hispidae | *Aspidimorpha furcata* (Thunberg, 1789) | Omnivore | | 1 | | 0.0070 |
| Staphylinidae | *Bolitogyrus fukienensis* (Scheerpeltz, 1974) | Omnivore | | 2 | | 0.0022 |
|  | *Paederus fuscipes* Curtis, 1826 | Omnivore | | 28 | | 0.1473 |
|  | *Paederus sondaicus* Fauvel, 1895 | Omnivore | | 2 | | 0.0012 |
| Latridiidae | *Lathridius minutus* L., 1767 | Saprophage | | 220 | | 0.0812 |
|  | *Migneauxia lederi* Reitter, 1875 | Saprophage | | 341 | | 0.2073 |
| Mycetophagidae | *Typhaea stercorea* (L., 1758) | Saprophage | | 1 | | 0.0004 |
| Silvanidae | *Ahasverus advena* (Waltl, 1834) | Saprophage | | 2 | | 0.0005 |
|  | *Psammoecus triguttatus* Reitter, 1874 | Saprophage | | 50 | | 0.0359 |
|  | *Silvanophus longicollis* (Reitter, 1876) | Saprophage | | 73 | | 0.0317 |
| Total | | | 28394 | | 1015.9060 | |
