## Supplementary Material 2 for "Effects of cover crops on beetle assemblages in tea plantations"

| Level 1 | Level 2 | Estimate | Std. Error | Df | Statistic | P-Value |
| --- | --- | --- | --- | --- | --- | --- |
| Pooled log-abundance lsmeans | | | | | | |
| Nat | Bar | -0.175 | 0.088 | 217 | -1.983 | 0.198 |
| Nat | Cas | -0.288 | 0.088 | 217 | -3.273 | 0.007 |
| Nat | Pas | -0.148 | 0.088 | 217 | -1.683 | 0.336 |
| Bar | Cas | -0.114 | 0.088 | 217 | -1.290 | 0.570 |
| Bar | Pas | 0.026 | 0.088 | 217 | 0.300 | 0.991 |
| Cas | Pas | 0.140 | 0.088 | 217 | 1.590 | 0.386 |
| Hervibore log-abundance lsmeans | | | | | | |
| Nat | Bar | -0.178 | 0.112 | 217 | -1.596 | 0.383 |
| Nat | Cas | -0.062 | 0.112 | 217 | -0.555 | 0.945 |
| Nat | Pas | 0.038 | 0.112 | 217 | 0.341 | 0.986 |
| Bar | Cas | 0.116 | 0.112 | 217 | 1.041 | 0.725 |
| Bar | Pas | 0.216 | 0.112 | 217 | 1.937 | 0.216 |
| Cas | Pas | 0.100 | 0.112 | 217 | 0.896 | 0.807 |
| Predator log-abundance lsmeans | | | | | | |
| Nat | Bar | -0.106 | 0.100 | 217 | -1.058 | 0.715 |
| Nat | Cas | -0.407 | 0.100 | 217 | -4.067 | 0.000 |
| Nat | Pas | -0.318 | 0.100 | 217 | -3.173 | 0.009 |
| Bar | Cas | -0.301 | 0.100 | 217 | -3.009 | 0.015 |
| Bar | Pas | -0.212 | 0.100 | 217 | -2.115 | 0.152 |
| Cas | Pas | 0.090 | 0.100 | 217 | 0.894 | 0.808 |
| Predator-herbivore log-abundance ratios lsmeans | | | | | | |
| Nat | Bar | 0.041 | 0.068 | 217 | 0.607 | 0.930 |
| Nat | Cas | 0.012 | 0.068 | 217 | 0.172 | 0.998 |
| Nat | Pas | -0.059 | 0.068 | 217 | -0.872 | 0.820 |
| Bar | Cas | -0.030 | 0.068 | 217 | -0.435 | 0.972 |
| Bar | Pas | -0.101 | 0.068 | 217 | -1.479 | 0.452 |
| Cas | Pas | -0.071 | 0.068 | 217 | -1.044 | 0.724 |
| Pooled log-biomass lsmeans | | | | | | |
| Nat | Bar | -0.196 | 0.148 | 217 | -1.324 | 0.548 |
| Nat | Cas | -0.170 | 0.148 | 217 | -1.148 | 0.660 |
| Nat | Pas | -0.097 | 0.148 | 217 | -0.653 | 0.914 |
| Bar | Cas | 0.026 | 0.148 | 217 | 0.176 | 0.998 |
| Bar | Pas | 0.100 | 0.148 | 217 | 0.672 | 0.908 |
| Cas | Pas | 0.073 | 0.148 | 217 | 0.495 | 0.960 |
| Herbivore log-biomass lsmeans | | | | | | |
| Nat | Bar | -0.204 | 0.170 | 217 | -1.201 | 0.626 |
| Nat | Cas | -0.122 | 0.170 | 217 | -0.717 | 0.890 |
| Nat | Pas | -0.092 | 0.170 | 217 | -0.545 | 0.948 |
| Bar | Cas | 0.082 | 0.170 | 217 | 0.485 | 0.962 |
| Bar | Pas | 0.112 | 0.170 | 217 | 0.656 | 0.913 |
| Cas | Pas | 0.029 | 0.170 | 217 | 0.172 | 0.998 |
| Predator log-biomass lsmeans | | | | | | |
| Nat | Bar | -0.124 | 0.083 | 217 | -1.496 | 0.442 |
| Nat | Cas | -0.260 | 0.083 | 217 | -3.13 | 0.010 |
| Nat | Pas | -0.242 | 0.083 | 217 | -2.913 | 0.020 |
| Bar | Cas | -0.136 | 0.083 | 217 | -1.642 | 0.357 |
| Bar | Pas | -0.118 | 0.083 | 217 | -1.417 | 0.490 |
| Cas | Pas | 0.019 | 0.083 | 217 | 0.225 | 0.996 |
| Predator-herbivore log-biomass ratios lsmeans | | | | | | |
| Nat | Bar | 0.027 | 0.064 | 217 | 0.419 | 0.975 |
| Nat | Cas | -0.046 | 0.064 | 217 | -0.721 | 0.888 |
| Nat | Pas | -0.049 | 0.064 | 217 | -0.777 | 0.865 |
| Bar | Cas | -0.073 | 0.064 | 217 | -1.141 | 0.665 |
| Bar | Pas | -0.076 | 0.064 | 217 | -1.196 | 0.630 |
| Cas | Pas | -0.004 | 0.064 | 217 | -0.055 | 1.000 |
| Pooled species richness lsmeans | | | | | | |
| Nat | Bar | -1.014 | 0.334 | 217 | -3.039 | 0.014 |
| Nat | Cas | -1.153 | 0.334 | 217 | -3.456 | 0.004 |
| Nat | Pas | -0.861 | 0.334 | 217 | -2.581 | 0.051 |
| Bar | Cas | -0.139 | 0.334 | 217 | -0.416 | 0.976 |
| Bar | Pas | 0.153 | 0.334 | 217 | 0.458 | 0.968 |
| Cas | Pas | 0.292 | 0.334 | 217 | 0.874 | 0.818 |
| Herbivore species richness lsmeans | | | | | | |
| Nat | Bar | -0.264 | 0.203 | 217 | -1.302 | 0.563 |
| Nat | Cas | -0.208 | 0.203 | 217 | -1.028 | 0.733 |
| Nat | Pas | -0.333 | 0.203 | 217 | -1.645 | 0.356 |
| Bar | Cas | 0.0556 | 0.203 | 217 | 0.274 | 0.993 |
| Bar | Pas | -0.069 | 0.203 | 217 | -0.343 | 0.986 |
| Cas | Pas | -0.125 | 0.203 | 217 | -0.617 | 0.927 |
| Predator species richness lsmeans | | | | | | |
| Nat | Bar | -0.417 | 0.171 | 217 | -2.441 | 0.072 |
| Nat | Cas | -0.653 | 0.171 | 217 | -3.824 | 0.001 |
| Nat | Pas | -0.347 | 0.171 | 217 | -2.034 | 0.179 |
| Bar | Cas | -0.236 | 0.171 | 217 | -1.383 | 0.511 |
| Bar | Pas | 0.069 | 0.171 | 217 | 0.407 | 0.977 |
| Cas | Pas | 0.306 | 0.171 | 217 | 1.790 | 0.281 |
| Predator-herbivore species richness ratios lsmeans | | | | | | |
| Nat | Bar | -0.016 | 0.028 | 217 | -0.559 | 0.944 |
| Nat | Cas | -0.034 | 0.028 | 217 | -1.207 | 0.623 |
| Nat | Pas | -0.007 | 0.028 | 217 | -0.258 | 0.994 |
| Bar | Cas | -0.018 | 0.028 | 217 | -0.648 | 0.916 |
| Bar | Pas | 0.009 | 0.028 | 217 | 0.301 | 0.990 |
| Cas | Pas | 0.027 | 0.028 | 217 | 0.949 | 0.778 |
