## Supplementary figures and images for "Effects of cover crops on beetle assemblages in tea plantations"

### Supplementary Material 3

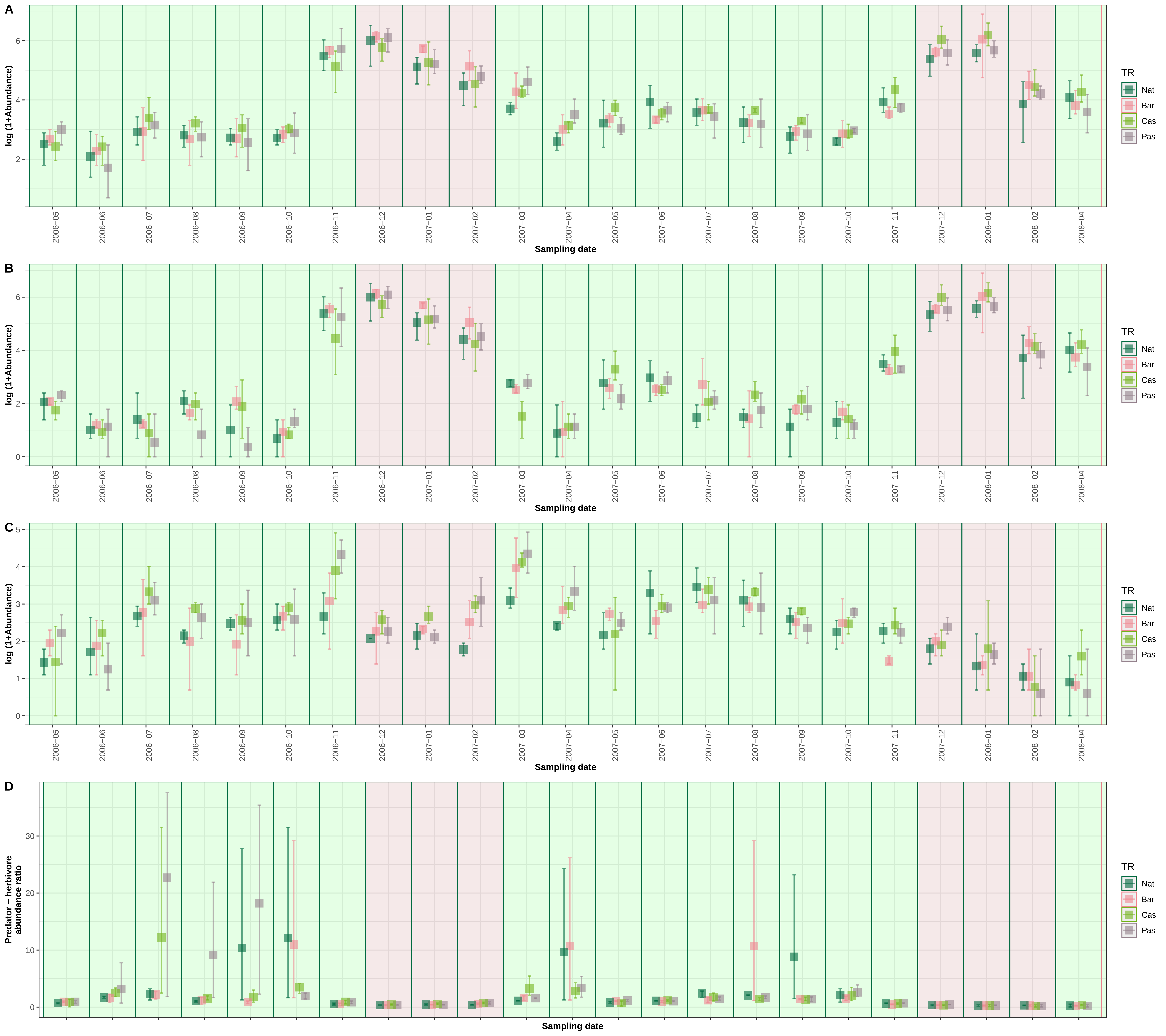

### Supplementary Material 4

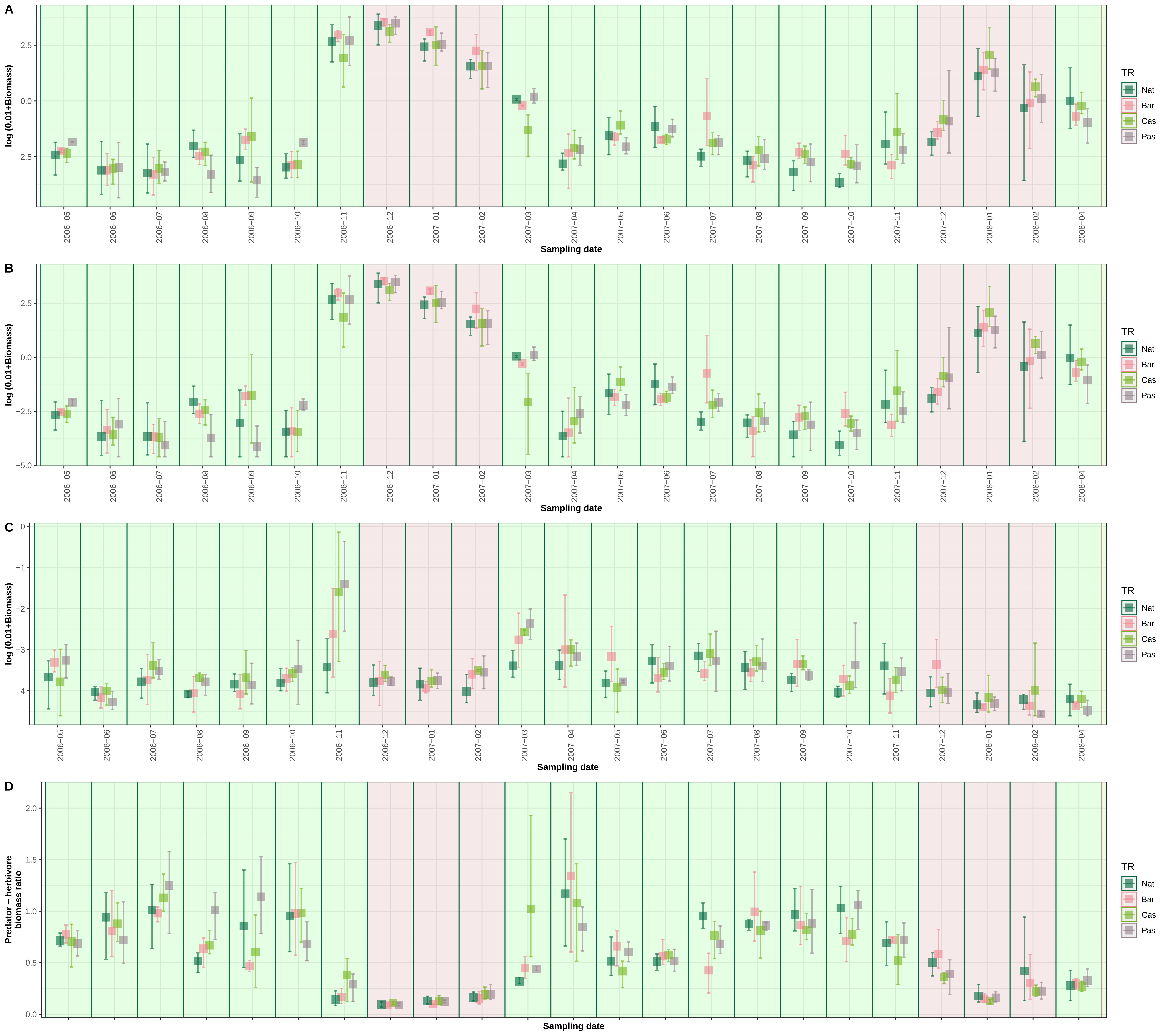

### Supplementary Material 5

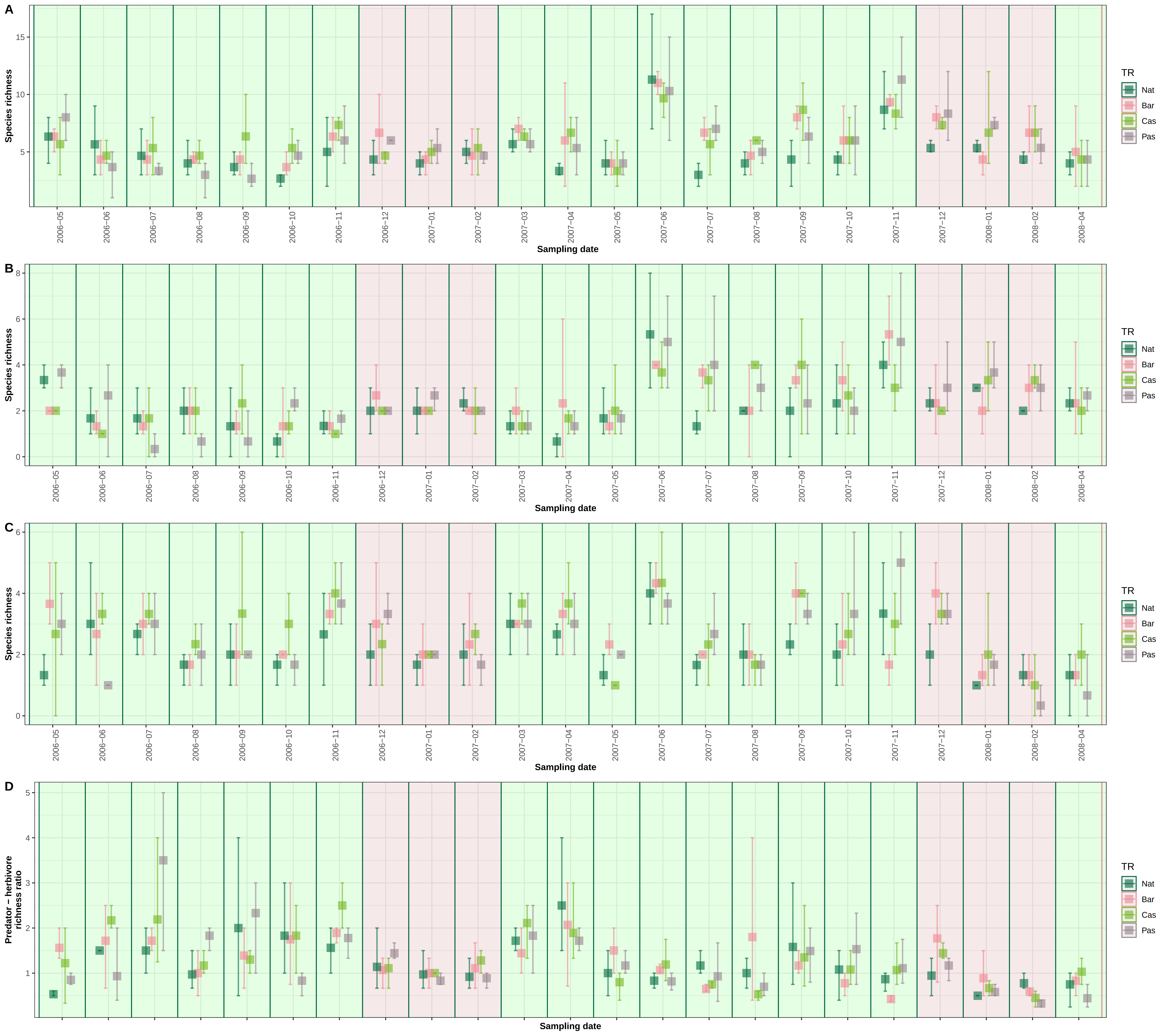
